# Intraspecific density changes the impact of interspecific competitors on parasite infection traits in multiple infections

**DOI:** 10.1101/2023.12.19.572385

**Authors:** Diogo P. Godinho, Leonor R. Rodrigues, Sophie Lefèvre, Sara Magalhães, Alison B. Duncan

## Abstract

Co-infections are frequent, with consequences for parasite life-history traits expressed at within- and between-host levels. However, little is known about whether the effect of interspecific competitors on traits are correlated or independent or if they change with intraspecific competition. To address this, we investigated the occurrence of genetic correlations among within- and between-host traits at different intra-specific densities of inbred lines of the spider mite, *Tetranychus urticae* with its competitor *T. evansi*. First, we found *T. evansi* presence on a shared host leaf produced a negative (non-genetic) correlation between virulence (leaf damage) and number of daughters (transmitting stages) at intermediate intraspecific densities; this same relationship was not significant without competitors. Second, we show interspecific competitors increases transmission to adjacent leaf discs, measured as movement of adult *T. urticae* females, but only at low and intermediate intraspecific densities. Finally we tested whether within-host traits (virulence and transmitting stages) were correlated with between-host traits (movement to adjacent patches), with or without competitors, at different conspecific densities. We found traits were mostly independent; interspecific competitors may increase transmission across hosts without affecting virulence (or vice versa). These independent effects on within- and between-host traits indicate competition may impact epidemiology and parasite trait evolution.

## Introduction

Interspecific interactions among different parasite species can affect the expression of several traits in parasites. Indeed, interspecific competitors can both increase or decrease within-host parasite growth e.g. (1-4) and virulence under multiple infections may differ from that under single infections (5-7). Finally interspecific interactions may also directly affect transmission by triggering dispersal from hosts infected with competitors or impacting whether a new host becomes infected. Indeed, certain parasites avoid or choose a host depending on its infection status (8, 9).

The effect of inter and intra-specific competition in multiple infection scenarios may depend on the relative densities of each competitor (2, 4, 10-12). Higher doses of an interspecific competitor may increasingly reduce a focal parasite’s growth (4, 11). Alternatively, higher intraspecific densities (competition) may change the impact of an interspecific competitor on parasite growth or host traits (2, 10, 12), even to the extent that the impact of interspecific competition is no longer apparent at certain (higher) intraspecific densities (10). The relative densities of conspecifics versus interspecific competitors may also affect parasite traits in the between host environment such as whether a parasite establishes on an already infected host (13) and/or movement between infected hosts.

Another important factor is that parasite traits are not necessarily independent. Indeed, the prediction that too high virulence leads to rapid host death curtailing transmission forms the basis of the trade-off hypothesis (14, 15). Thus, it is important to consider multiple infection scenarios in the study of virulence-transmission relationships because competitors may change outcomes (16-18). In this sense, it is key to disentangle the relative impact of interspecific interactions on within-or between host parasite traits and to test whether such effects are direct or indirect and genetic or environmental. To our knowledge, this has not been done.

This study uses isogenic lines of the spider mite *Tetranychus urticae* to investigate how the presence of an interspecific competitor, the closely related species *T. evansi*, impacts virulence, the number of adult daughters and transmission (i.e.,. time to reach, and numbers reaching, a new host patch) as well as the potential (genetic) correlations among these traits. In this species, there is genetic variation for dispersal distance (19, 20) (which we refer to, and is the same as, transmission) and the propensity for aerial dispersal (21). Moreover, dispersal is a plastic trait, with individuals having higher dispersal at elevated intraspecific densities (19) and in the presence of kin (22). Previous work has investigated how dispersal is linked to other life-history traits in *T. urticae*. For example, selection for higher dispersal has been shown to be associated with higher diapause incidence and lower fecundity (20), though this may be due to persisting epigenetic effects (23). There are also phenotypic relationships among traits whereby dispersing individuals have fewer descendants when laying at higher densities (24) and have smaller eggs (25). Further, in a recent companion study to this one with the same inbred lines, we found a positive genetic correlation between virulence and number of adult daughters when transmission was possible during the infectious period (26). Hence, there are both genetic and environmental relationships between transmission and other life-history traits.

On tomato plants, *T. evansi* generally outcompetes *T. urticae* (12, 27) but see (28). However, outcomes can change when between-host dynamics determine within-host dynamics, for example when the sequence of arrival determines the outcome of competition. For instance, Fragata et al (2022) showed that *T. evansi* excludes *T. urticae* except when the latter arrived first and occupied *T. evansi’s* preferred niche (13). That order of arrival in multiple infections changes what happens in the within-host environment is consistent with theoretical predictions (16-18).

We used inbred lines of *T. urticae* to investigate how interspecific competitors affect the relationship between virulence and the number of adult females produced, as well as transmission, at different intraspecific densities. Moreover, we evaluate whether traits measured within hosts correlate with those expressed between hosts, in the presence or absence of competitors.

## Materials and Methods

### Biological system

Spider mites are macro-parasites of plants, including many economically important crops, with their complete life cycle occurring on their host plant (29). Both *Tetranychus urticae* and *T. evansi* females lay eggs on leaves, which take ∼4 days to hatch. The juvenile stage comprises 1 nymph stage and 2 deuteronymph stages, with adults emerging approximately after 14 days in our laboratory. All stages feed by injecting their stylet into parenchyma cells and sucking out the cytoplasm, which leaves chlorotic damage on the leaf surface, our measure of virulence (30). *T. urticae* is a generalist species, feeding on more than 1000 different plant species (31), whereas *T. evansi* is a specialist species, mostly feeding on *Solanaceae* plants (32). In natural systems, co-occurrence of difference spider mite species in the same geographical area is common, leading to co-infection of the same host plant (33).

### Spider-mite populations

Inbred lines of *Tetranychus urticae* were created from an outbred population through 14 generations of sib mating at the University of Lisbon (33). A subset of 15 inbred lines was transferred to the University of Montpellier in January 2018 and maintained on bean leaves (variety Pongo) as described in Godinho et al. 2023. The *Tetranychus evansi* population was collected in October 2010 in the Alpes Maritimes (43.75313 N, 7.41977 E) on *Solanum nigrum*. A subset was transferred to the University of Montpellier in January 2018, where they were maintained on bean leaves (variety Pongo).

Prior to each experiment, cohorts of 40 mated female spider mites from each inbred line were isolated on a bean patch (2-3 leaves placed together). These females were allowed to lay eggs for 48h. Fourteen days later, the mated daughters of these females, of approximately the same age, were used in the experiments. The same procedure was used to create cohorts of *T. evansi*. All spider-mite populations, inbred lines and cohorts used in these experiments were maintained on bean leaves (variety Pongo) placed on water saturated cotton wool, in small plastic boxes (255 mm length x 183 mm width x 77 mm height), at 25°C with a 16:8 L: D cycle, at 60% relative humidity. Not all inbred lines are represented in each experiment due to too few individuals available at the start of the experiment (N between 12 to 14 lines).

#### 1. The impact of interspecific competitors on within-host traits and their correlation

Females of each *T. urticae* inbred lines were randomly assigned to one ‘intraspecific density’ treatment (5, 10 and 20 females), with or without ‘interspecific competition’ (10 *T. evansi* females). In all treatments, females were placed on a 2 × 2 cm bean leaf patch placed on wet cotton wool in plastic boxes (Figure S1a). There were 5 to 13 replicates for each inbred line per treatment combination (intraspecific density x interspecific competition) distributed across 3 blocks. All females were allowed to feed and lay eggs on their leaf patches for 4 days. After this period, females were killed, the number of eggs was counted and a photograph of each patch taken using a Canon EOS 70D camera. The damage inflicted by the spider mites, used as a measure of virulence, was determined using imageJ and Ilastik 1.3, as described in (26). Succinctly, the background from each photo was removed in imageJ, subsequently using Ilastik we distinguished damaged area from healthy leaf and then in imageJ the damaged area was calculated via the colour contrast between damaged and undamaged leaf tissue. Because some leaf veins were assigned as damage by Ilastik, uninfested bean leaf patches were left in the experimental boxes for the same period of time and photographed. These control patches were used to establish an average baseline level of damage, which was subtracted from each measurement. After a period of 14 days, the female offspring surviving on each patch was counted. Only females were counted because the males of both species are not easily distinguishable, females are the main dispersers in these species and the number of females produced correlates with transmission (Godinho et al. 2023). As previously stated, the data on damage inflicted and production of adult females, for the “intraspecific density” treatments in the absence of *T. evansi* is published elsewhere (26).

#### 2. The effects of interspecific competition on transmission

We measured differences in dispersal (between-host dynamics) for the different *T. urticae* inbred lines assigned to the same ‘intraspecific density’ and ‘interspecific competition’ treatments as in Experiment 1. Adult *T. urticae* females were placed in groups of 5, 10 or 20 on a 2cm^2^ bean leaf patch on wet cotton wool alone or with 10 *T. evansi* females. This first host patch was connected, in a row, to 2 other bean leaf host patches via 3 × 1 cm Parafilm bridges from day 1 of the experiment (Figure S1b). This experimental setup was replicated across several boxes. Females were allowed to feed and disperse across the patches, and the number of mites on each patch counted on days 1, 2, 3, 6, and 9 after the beginning of the experiment. There were 3 to 13 replicates for each inbred line per treatment combination (intraspecific density x interspecific competition) distributed across 2 blocks.

### Statistical analysis

Analyses were preformed using the software JMP SAS version 17 and SAS OnDemand for Academics (34).

#### 1. Impact of interspecific competitors on within-host traits

In Experiment 1 General Linear Mixed Models (GLMM) were used to investigate how intraspecific competition (density) and interspecific competition affected virulence and the number of female offspring becoming adult. Analyses for experiment 1 included intraspecific density as a covariate and interspecific competition as a fixed factor. When a term including intraspecific or interspecific competition in the model was significant, post-hoc Tukey tests were used to look for differences among treatments.

#### 2. Impact of interspecific competitors on between-host traits

In Experiment 2, different measures were taken to assess dispersal across host patches. We used a Dispersal score to evaluate the spread of mites across the 3 host patch system. This was calculated as the ((number of mites on host patch 2) + (the number of mites on host patch 3*2))/total number of mites) on each day measurements were taken (19). The dispersal score was analysed in a GLMM with interspecific competition included in the model as a fixed factor, intraspecific density and time as covariates and their interactions. We also investigated, in separate GLMMs, including intraspecific density as a covariate and interspecific competition as a fixed factor, how interspecific competition affected the time for mites to reach and the maximum number of *T. urticae* on host patches 2 and 3. Full models included interactions between explanatory variables which were simplified removing nonsignificant terms in a stepwise fashion. All the above models included inbred line and block as random factors.

#### Broad sense heritability

Broad-sense heritability, 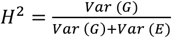 for each trait in each experiment was determined by extracting the proportion of total variance in models explained by inbred line (among inbred line variance) by re-running models for dispersal related traits including all terms as random (competition, line, block and/or density) to obtain all variance components. Note, models did not include interactions between terms. Traits, when possible, were divided by the total number of adult females placed on a patch because heritability is an individual trait. The significance of each model was assessed by comparing the Akaike’s Information Criterion (AIC) of models including inbred line with a model excluding it.

#### Correlations between traits measured in the within and between host environments

We assessed genetic correlations measured between traits in the 2 experimental set-ups separately for each combination of density and competition treatments. In order to randomly couple, multiple times, measures in the different experiments we bootstrapped (with replacement) the mean value for each inbred line at each density and interspecific competition treatment 20 times. The actual mean values for each inbred line for pairs of traits were coupled and the 20 bootstrapped means randomly coupled. Mean values were standardised across all lines for each density and competition treatment so each variable had a mean of zero and standard deviation of one. This means values of pairs of traits were of a similar scale as required for the PROC MIXED COVTEST in SAS Studio.

Genetic correlations were done using a PROC MIXED COVTEST model as described in (36) using SAS Studio. Briefly, each paired value was given an identity and the columns for each pair of traits stacked giving 2 columns, one showing the 2 traits measured (labelled as trait) and the other the values (the response variable). Trait was included in the model as an explanatory variable. There were 2 random terms in the model. The first random term included an interaction between ‘trait’ and inbred line, thus assessing the among inbred line variance for each trait. The error structure for this term was specified as ‘unr’, which tells the model to assess the genetic correlation across the 2 traits. A second random term was included in the model with the trait pair ‘ID’ nested within inbred line. The significance of the genetic correlation from each model was given using a log-likelihood ratio test by comparing the log likelihood of the aforementioned model with a model where the error structure was defined as ‘un(1)’, thus constraining the covariance matrix to zero.

Phenotypic (Spearmans) correlations were done between virulence and the number of adult daughters in Experiment 1 across the standardised measures within each density and competition treatment combination. As it only made sense to look at phenotypic correlations when traits were measured in the same experimental set up, this was only done for this pair of traits. All p-values < 0.05 were corrected for multiple testing (within each pair of traits) using the Bonferroni correction method. The dispersal score was not included as a trait in the correlations as it was measured through time, making it difficult to obtain a single measure. As measures were taken across different experiments, block was not included in these analyses.

## Results

### 1. The impact of interspecific competitors on within-host traits and their correlation

The presence of interspecific competition at intermediate conspecific density made the phenotypic correlation between virulence and the number of daughters negative (Figure 1b; Table S1), while in the absence of competition there was no correlation. At low and high conspecific densities, both in the presence and absence of *T. evansi*, the phenotypic correlations between these traits were positive and negative respectively (Figure 1; Table S1), and there were no genetic correlations between traits (Table S1), corroborating the results found in Godinho et al (2023) (26).

**Figure 1:**
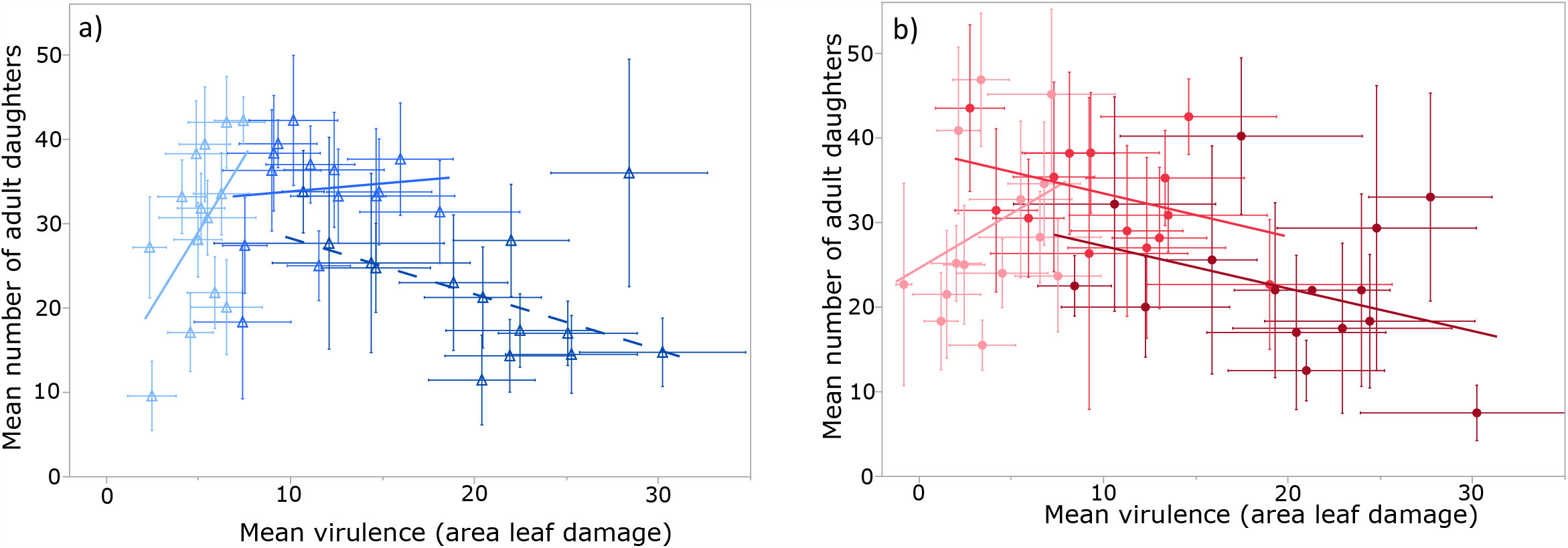
shows the relationship between virulence and transmission/dispersal potential at low, intermediate and high density in a) the absence (blue) and b) presence (red) of 10 *T. evansi* competitors. Mean values at low densities are lighter in colour, at intermediate densities medium colour and high densities darker colours. Each point is the mean value for an inbred line at each density (+ SE).

The presence of *T. evansi* had no effect on *T. urticae* virulence or the number of adult daughters (Figure 1 and S2; Table S2; no significant main effect of, or interaction terms including ‘interspecific competition’). As these results were not affected by interspecific competition they are not discussed further, but are presented elsewhere (26).

### 2. Impact of interspecific competitors on between-host traits

The dispersal score was affected by both interspecific competition and intraspecific density, with a significant interaction between these two factors (Figure 2, Table S3). This interaction showed that, at low and intermediate density,*T. urticae* females were more likely to leave the patch earlier in the presence of *T. evansi*. The interaction between *T. urticae* density and time was also significant, with values of the dispersal score saturating through time for patches in the intermediate and high density treatments (Figure 1, Table S3). The dispersal score captured time to arrival on, and the maximum number of, spider mites on host patches 2 and 3 (Figure S3, Table S4).

**Figure 2:**
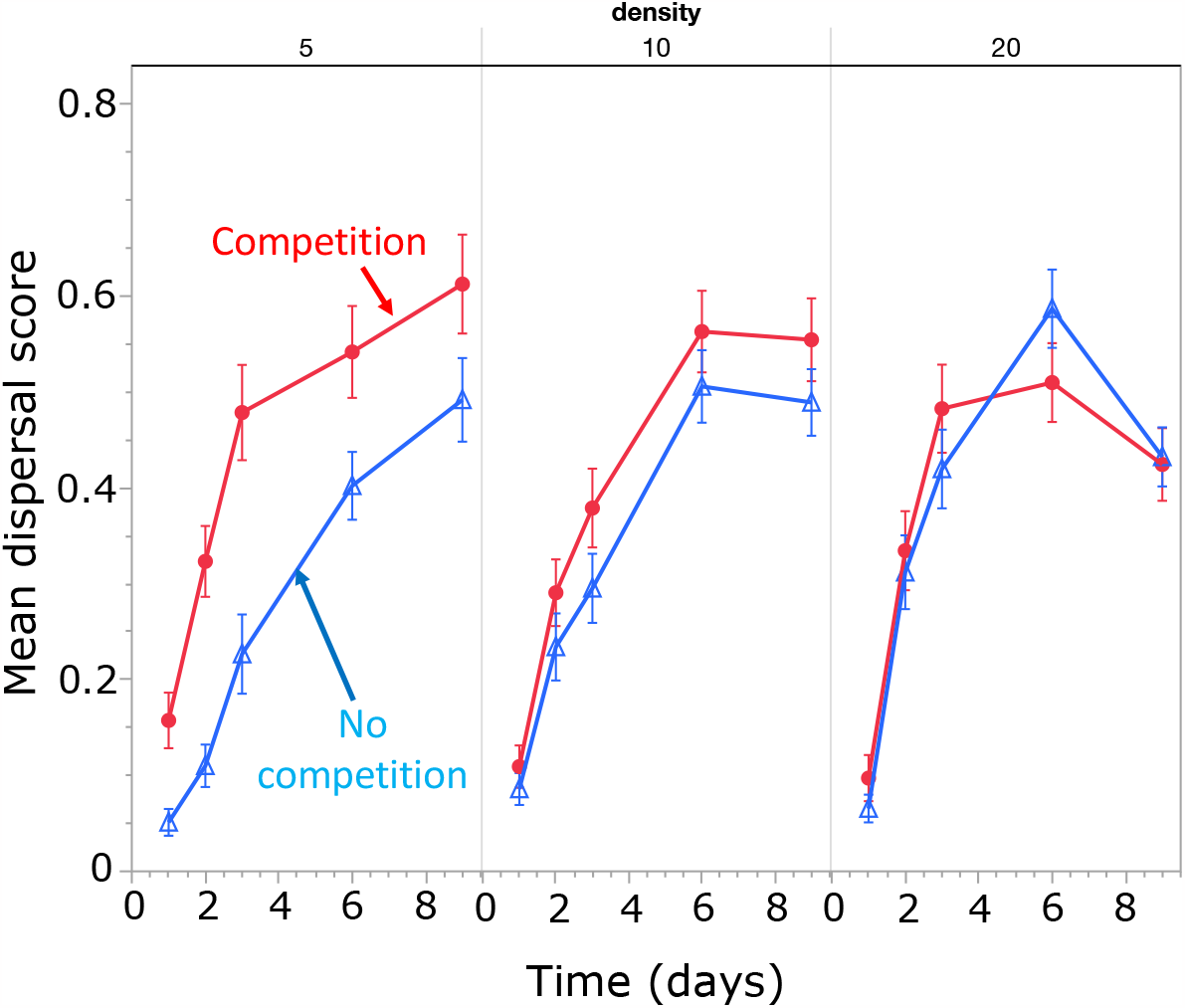
dispersal score (number of mites on host patch 2 × 1 + number of mites on host patch 3 × 2)/density) at each of the different *T. urticae* densities in the presence (red) or absence (blue) of competition with *T. evansi*.

### Genetic variance for within-host dynamics and between-host dynamics related traits

In the within-host environment, the number of adult daughters had a genetic component (H2 = 0.057). In the between host environment, inbred line explained a significant portion of the variance (H2) for time to reach host patches 2 and 3, the maximum number on patch 3 and the dispersal score, but not maximum number on patch 2 (Table S5).

### Genetic correlations between traits measured in the within and between-host environment

We aimed to explore the genetic relationships between 8 traits in 6 different treatment combinations of intraspecific density and interspecific competition giving a total of 48 correlations. Of these, 9 models did not converge, likely due to a lack of genetic variation for both or one of the traits at higher densities (or power due to fewer replicates). Note that for all models that converged inbred line explained a significant portion of the variance. This may differ from H2 calculations since these analyses were done on bootstrapped data. Of the remaining 39 models investigating genetic correlations between traits, only 2 were significant. This shows that virulence and number of adult daughters measured in the within-host environment are mostly independent of traits measured in the between host environment (Table 1). The 2 significant correlations show that 1. there was a negative correlation between virulence and the time to arrive on host patch 2 at high *T. urticae* densities in the presence of interspecific competition (Figure S4a) and 2. at intermediate *T. urticae* density, in the absence of interspecific competition, there was a negative relationship between the number of adult daughters and the maximum number on host patch 2 (Figure S4b).

**Table 1:**
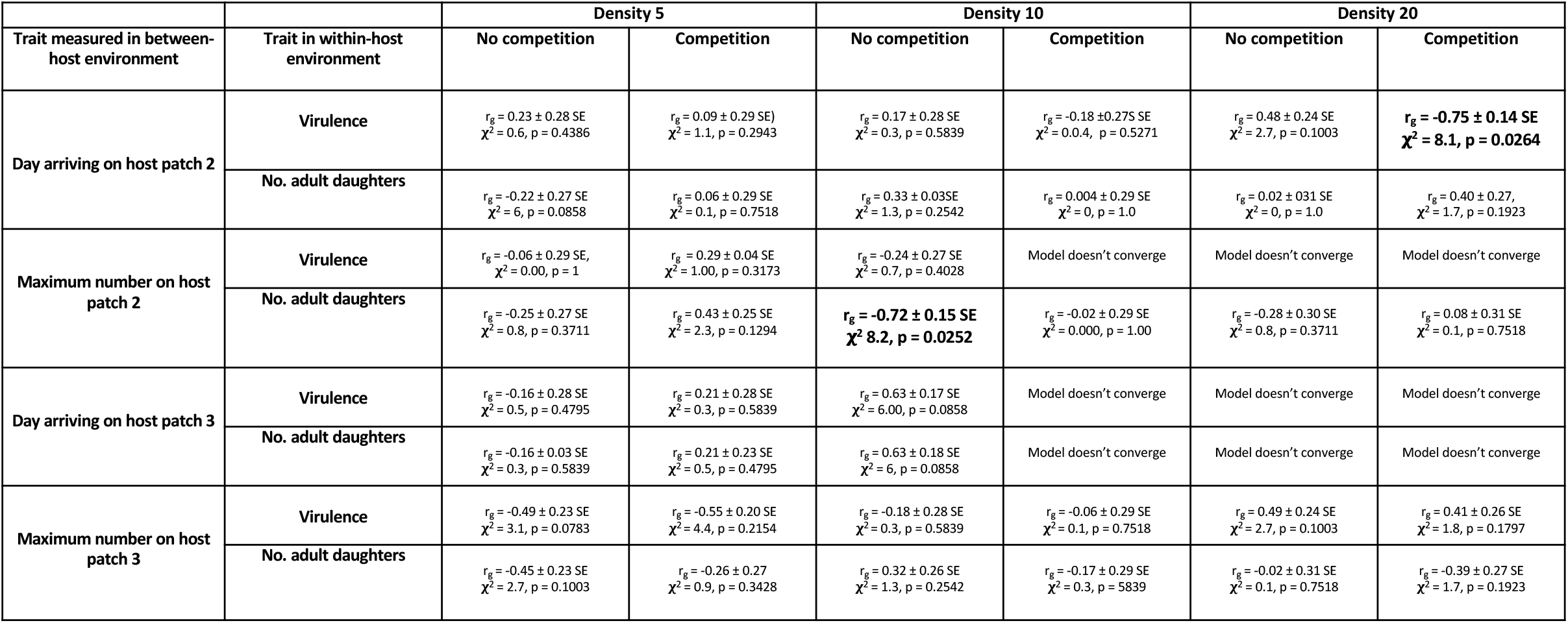
Summary of genetic correlations between transmission related traits measured in the between and within-host environment at each of the different intraspecific densities and in the presence and absence of interspecific competition. The genetic correlation (r_g_) for each pair of traits is presented ± the standard error and the result of the log likelihood test comparing models with and without the genetic correlation. All values of p < 0.05 were corrected using Bonferronni (counting 6 tests per pair of traits). Significant correlations are shown in bold.

## Discussion

We found that the presence of interspecific competitors caused the relationship between virulence and the number of daughters produced to become negative at lower intraspecific densities. The presence of interspecific competitors also caused transmission of *T. urticae* to new host patches sooner at low and intermediate intraspecific densities. This, for the most part, was genetically unrelated to measures of virulence or number of adult daughters in the within-host environment. This was despite many traits in both the within and between host environment being genetically determined. Thus, selection for virulence or the number of adult daughters in the within-host environment is mostly unlinked to traits describing between-host transmission. This means that selection for faster spread across host patches is not necessarily associated with higher virulence.

### The impact of interspecific competitors on within-host traits and their correlation

The presence of interspecific competitors did not affect levels of virulence or the production of transmitting stages per se. It did however, cause the relationship between these traits to become negative at intermediate intraspecific densities. In the absence of interspecific competitors there is only a negative relationship between these traits at high densities. These results show that the additional impact of *T. evansi* depends on the intensity of intraspecific competition. A number of studies have shown that the impact of interspecific competition on transmission-related traits can change in response to the relative densities of parasites in coinfection (2, 4, 11). These studies generally show that, if that the negative affect of an interspecific competitor increases with its infection dose, and presumably density (2, 4). However, Fellous et al (2009) found, as we do, that the impact of interspecific competition on parasite growth also depends on intraspecific parasite densities (2).

It should be noted that these results may have been very different had this experiment been done on a different host plant to which *T. evansi* is better adapted. Indeed, *T. evansi* is generally found to be the superior competitor on tomato plants, often excluding *T. urticae* (12, 27) but see (28). Nevertheless, Fragata et al (2022) found that these species can co-exist if *T. urticae* arrives first on tomato and occupies *T. evansi’s* preferred niche (13), suggesting, thus, that it can also depend on other factors in the environment such as the overall prevalence of both species.

### Interspecific competition affected between-host traits

Interspecific competition did change how *T. urticae* moved among host patches, but as for in the within-host environment, effects depended on the intensity of intraspecific competition. At low and intermediate intraspecific densities, the presence of *T. evansi* caused *T. urticae* females to move to the second and third host patch sooner and to be on these host patches at higher densities. At high *T. urticae* densities, however, interspecific competitors did not affect transmission. This is probably because the density of *T. urticae* was so high that there was no additional effect of interspecific competition.

The finding that interspecific competition causes *T. urticae* to move to a new host faster means that coinfection may be an important driver of epidemic spread. Coinfection can cause individuals to become superspreaders, when an infected host is responsible for a disproportionate number of transmission events (37). Here we only measured the number of spider mites moving from one host-patch to another, which is not the same as the number of new hosts infected. Nevertheless, it gives an idea of the number of transmission stages leaving an infected host which is a measure of infection potential, similar to parasite shedding (38-41). These different effects of parasite intraspecific densities and coinfection could be used as a predictor of parasite spread in natural populations and used to manage or control epidemics, such as identifying (and isolating or treating) the most infectious individuals (42).

### Limited interactions among within- and between-host traits

Our results show that within-host traits are mostly genetically independent of the among host traits measured. There was no genetic relationship among traits in 46/48 possible tests across treatment combinations, despite these traits being genetically determined under many conditions. This contrasts with our finding of a positive genetic correlation between virulence and transmission, in the absence of interspecific competition, when transmission is from the same patch where virulence is measured (26). Hence, it may be possible to relate higher virulence to the number of transmission stages produced from that host, but this does not necessarily tell us how these transmission stages will behave independently of within-host processes. A line with higher virulence, or that produces more offspring, is not necessarily going to move more quickly across host patches.

How such direct measures of transmission in the between host environment, independent of withinhost processes, actually scale up and affect the spread of *T. urticae* across a population of potential hosts is not clear. Different life-history strategies could co-exist in a parasite population, some maximising fitness in the within-host environment with others spreading faster across the host population. If genetic variation for within and between-host traits are uncoupled, then contrasting selection pressures in each environment may maintain variation for both across scales. Alternatively, if density dependence had been less intense in the within-host environment we may have found genetic correlations among traits. Indeed, in a previous study when transmission was possible from the same leaf patch virulence was measured, thus alleviating intense within-host density dependence, there was a positive genetic correlation between virulence and transmission (26).

Only 2 genetic correlations between traits measured in the within and between host environment were found and again these were dependent on the intensity of intraspecific competition and the presence of *T. evansi*. In the absence of interspecific competition, at intermediate *T. urticae* densities, there was a negative relationship between mean number of adult daughters and the maximum number of females on the second host patch. It is possible that different strategies exist whereby lines that are not so competitive in the within-host environment, are more likely to move and establish on a new host, consistent with the competition-colonisation trade-off (43, 44). The fact that this was not observed in the presence of interspecific competition may be because the presence of additional competition means these lines are not able to obtain sufficient resources to disperse. The second correlation was at high intraspecific densities, in the presence of *T. evansi*, there was a negative correlation between virulence and time to arrive on host patch 2. As this was dependent on the presence of interspecific competition, this means that these more virulent lines are responding to the presence of *T. evansi* as a trigger to leave the first host patch sooner.

### The impact of interspecific competition depends on the intensity of intraspecific competition

We find that the impact of interspecific competition in both the within and between host environment is only seen at lower intensities of intraspecific competition, as found in another study in the within-host environment looking at effects on host traits (10). A few studies have explored how the intensity of intraspecific parasite competition changes the impact of an interspecific competitor parasite in multipe infections (2, 4, 10, 12, 13), but results are not always presented as a function of changing intraspecific densities (e.g.(4)). In the within-host environment one study showed that the relative impact of interspecific competition on *T. urticae* growth declined with increasing *T. urticae* densities (12). Another found that the impact of an interspecific competitor (a helminth macroparasite) on shared host resources (red blood cells) was neglible at higher malaria densities, attributed to more intense intraspecifc malaria competition (10).

In the between-host environment it is less clear how the relative impact of inter and intraspecific competition among parasites play out. One study found order of arrival and the relative density affected probability of establishment and abundance in *T. evansi* and *T*.*urticae* (13). However, we are not aware of studies exploring how the interplay between inter and intraspecific parasite densities affects movement among hosts.

## Conclusion

Our results show that interspecific competition may increase the rate of parasite spread across hosts and that this trait is independent of traits measured in the within-host environment. This may mean that parasites selected for higher virulence locally are not those necessarily favoured in travelling wave epidemics, or those that spread far to seed infections in new host populations. Here, we only differentiate between indirect measures of transmission in the within-host environment and between host traits that are more direct measures of transmission. However, in future it would be interesting to explore how these traits are actually related to traits favouring the infection of a greater quantity of hosts or longer distant spread.

## Acknowledgements

We would like to thank Oliver Kaltz and the Experimental Evolution of Communities team at ISEM for helpful discussions. This is ISEM contribution number XXXX.

## References

1. Duncan AB, Agnew P, Noel V, Michalakis Y. The consequences of co-infections for parasite transmission in the mosquito Aedes aegypti. J Anim Ecol. 2015;84(2):498–508.

2. Fellous S, Koella JC. Infectious dose affects the outcome of the within-host competition between parasites. Am Nat. 2009;173(6):E177–84.

3. Hughes WO, Boomsma JJ. Let your enemy do the work: within-host interactions between two fungal parasites of leaf-cutting ants. Proc Biol Sci. 2004;271 Suppl 3:S104–6.

4. Ramsay C, Rohr JR. Identity and density of parasite exposures alter the outcome of coinfections: Implications for management. Journal of Applied Ecology. 2022;60(1):205–14.

5. Choisy M, de Roode JC. Mixed infections and the evolution of virulence: effects of resource competition, parasite plasticity, and impaired host immunity. Am Nat. 2010;175(5):E105–18.

6. Alizon S, de Roode JC, Michalakis Y. Multiple infections and the evolution of virulence. Ecol Lett. 2013;16(4):556–67.

7. Zele F, Magalhaes S, Kefi S, Duncan AB. Ecology and evolution of facilitation among symbionts. Nat Commun. 2018;9(1):4869.

8. Laidemitt MR, Gleichsner AM, Ingram CD, Gay SD, Reinhart EM, Mutuku MW, et al. Host preference of field-derived Schistosoma mansoni is influenced by snail host compatibility and infection status. Ecosphere. 2022;13(4).

9. Godinho DP, Janssen A, Li D, Cruz C, Magalhaes S. The distribution of herbivores between leaves matches their performance only in the absence of competitors. Ecol Evol. 2020;10(15):8405–15.

10. Wait LF, Kamiya T, Fairlie-Clarke KJ, Metcalf CJE, Graham AL, Mideo N. Differential drivers of intraspecific and interspecific competition during malaria-helminth co-infection. Parasitology. 2021;148(9):1030–9.

11. Ge S, Zheng D, Zhao Y, Liu H, Liu W, Sun Q, et al. Evaluating viral interference between Influenza virus and Newcastle disease virus using real-time reverse transcription–polymerase chain reaction in chicken eggs. Virology. 2012;9:1 –8.

12. Alzate A, Onstein RE, Etienne RS, Bonte D. The role of preadaptation, propagule pressure and competition in the colonization of new habitats. Oikos. 2020;129(6):820–9.

13. Fragata I, Costa-Pereira R, Kozak M, Majer A, Godoy O, Magalhaes S. Specific sequence of arrival promotes coexistence via spatial niche pre-emption by the weak competitor. Ecol Lett. 2022;25(7):1629–39.

14. Anderson RM, May RM. Coevolution of hosts and parasites. Parasitology. 1982;85 (Pt 2):411–26.

15. Alizon S, Hurford A, Mideo N, Van Baalen M. Virulence evolution and the trade-off hypothesis: history, current state of affairs and the future. J Evol Biol. 2009;22(2):245–59.

16. Mideo N, Alizon S, Day M. Linking within- and between-host dynamics in the evolutionary epidemiology of infectious diseases. Trends in Ecology and Evolution. 2008;23:511 –7.

17. Alizon S, van Baalen M. Multiple infections, immune dynamics, and the evolution of virulence. Am Nat. 2008;172(4):E150–68.

18. Coombs D, Gilchrist MA, Ball CL. Evaluating the importance of within- and between-host selection pressures on the evolution of chronic pathogens. Theor Popul Biol. 2007;72(4):576–91.

19. Bitume EV, Bonte D, Magalhaes S, San Martin G, Van Dongen S, Bach F, et al. Heritability and artificial selection on ambulatory dispersal distance in Tetranychus urticae: effects of density and maternal effects. PLoS One. 2011;6(10):e26927.

20. Yano S, Takafuji A. Variation in the life history pattern of Tetranychus urticae (Acari: Tetranychidae) after selection for dispersal. Experimental and Applied Acarology. 2002;27:1–10.

21. Li J, Margolies DC. Responses to direct and indirect selection on aerial dispersal behaviour in Tetranychus urticae. Heredity. 1994;74:10 –22.

22. Bitume EV, Bonte D, Ronce O, Bach F, Flaven E, Olivieri I, et al. Density and genetic relatedness increase dispersal distance in a subsocial organism. Ecol Lett. 2013;16(4):430–7.

23. Masier S, Bonte D. Spatial connectedness imposes local- and metapopulation-level selection on life history through feedbacks on demography. Ecology Letters. 2019;23(2):242–53.

24. Bonte D, De Roissart A, Wybouw N, Van Leeuwen T. Fitness maximization by dispersal: evidence from an invasion experiment. Ecology. 2014;95(11):3104–11.

25. Dahirel M, Masier S, Renault D, Bonte D. The distinct phenotypic signatures of dispersal and stress in an arthropod model: from physiology to life history. J Exp Biol. 2019;222(Pt 16).

26. Godinho DP, Rodrigues LR, Lefevre S, Delteil L, Mira AF, Fragata IR, et al. Limited host availability disrupts the genetic correlation between virulence and transmission. Evol Lett. 2023;7(1):58–66.

27. Sarmento RA, Lemos F, Dias CR, Kikuchi WT, Rodrigues JC, Pallini A, et al. A herbivorous mite down-regulates plant defence and produces web to exclude competitors. PLoS One. 2011;6(8):e23757.

28. Orsucci M, Navajas M, Fellous S. Genotype-specific interactions between parasitic arthropods. Heredity (Edinb). 2017;118(3):260–5.

29. Helle W, Sabelis MW. Spider Mites: Their Biology, Natural Enemies and Control. Amsterdam: Elsevier; 1985.

30. Mira AF, Marques L, Magalhães S, Rodrigues LR A method to measure the damage caused by cell-sucking herbivores.. In: Paula Duque DS, editor. Environmental Responses in Plants Methods and Protocols 2022. p. 299–312.

31. Grbic M, Van Leeuwen T, Clark RM, Rombauts S, Rouze P, Grbic V, et al. The genome of Tetranychus urticae reveals herbivorous pest adaptations. Nature. 2011;479(7374):487–92.

32. Navajas M, de Moraes GJ, Auger P, Migeon A. Review of the invasion of Tetranychus evansi: biology, colonization pathways, potential expansion and prospects for biological control. Exp Appl Acarol. 2013;59(1-2):43–65.

33. Godinho DP, Cruz MA, Charlery de la Masseliere M, Teodoro-Paulo J, Eira C, Fragata I, et al. Creating outbred and inbred populations in haplodiploids to measure adaptive responses in the laboratory. Ecol Evol. 2020;10(14):7291–305.

34. SAS OnDemand. Inc SI. Cary, NC, USA2023.

35. Falconer DS, Mackay TFC. Introduction to quantitative genetics: Longman; 1996.

36. Fry JD. Estimation of genetic variances and covariances by restricted maximum likelihood using PROC MIXED.

37. Lloyd-Smith J O., Schreiber SJ, Kopp PE, Getz WM. Superspreading and the effect of individual variation on disease emergence. Nature. 2005;435:355 –9.

38. Siva-Jothy JA, Vale PF. Dissecting genetic and sex-specific sources of host heterogeneity in pathogen shedding and spread. PLoS Pathog. 2021;17(1):e1009196.

39. Susi H, Barres B, Vale PF, Laine AL. Co-infection alters population dynamics of infectious disease. Nat Commun. 2015;6:5975.

40. Lass S, Hudson PJ, Thakar J, Saric J, Harvill E, Albert R, et al. Generating super-shedders: coinfection increases bacterial load and egg production of a gastrointestinal helminth. J R Soc Interface. 2013;10(80):20120588.

41. Lawley TD, Clare S, Walker AW, Goulding D, Stabler RA, Croucher N, et al. Antibiotic treatment of clostridium difficile carrier mice triggers a supershedder state, spore-mediated transmission, and severe disease in immunocompromised hosts. Infect Immun. 2009;77(9):3661–9.

42. Paull SH, Song S, McClure KM, Sackett LC, Kilpatrick AM, Johnson PT. From superspreaders to disease hotspots: linking transmission across hosts and space. Front Ecol Environ. 2012;10(2):75–82.

43. Nee S, May RM. Dynamics of metapopulations: habitat destruction and competitive coexistence.. Journal of Animal Ecology. 1992;61:37–40.

44. Hastings A. Coexistence of competitors in patchy environments. Ecology. 1980;75:40–7.

