## supplementary materials for "Intraspecific density changes the impact of interspecific competitors on parasite infection traits in multiple infections"

Figure S1: Experimental set up in a) Experiment 1 in which adult female *T. urticae* were placed in groups of 5, 10 or 20 on a $2cm^2$ bean leaf patch with or without 10 *T. evansi*, b) Experiment 2 in which adult female *T. urticae* were placed in groups of 5, 10 or 20 on a 3 x $2cm^2$ bean leaf patch system with or without 10 *T. evansi,* the picture here only depicts the low density treatment.

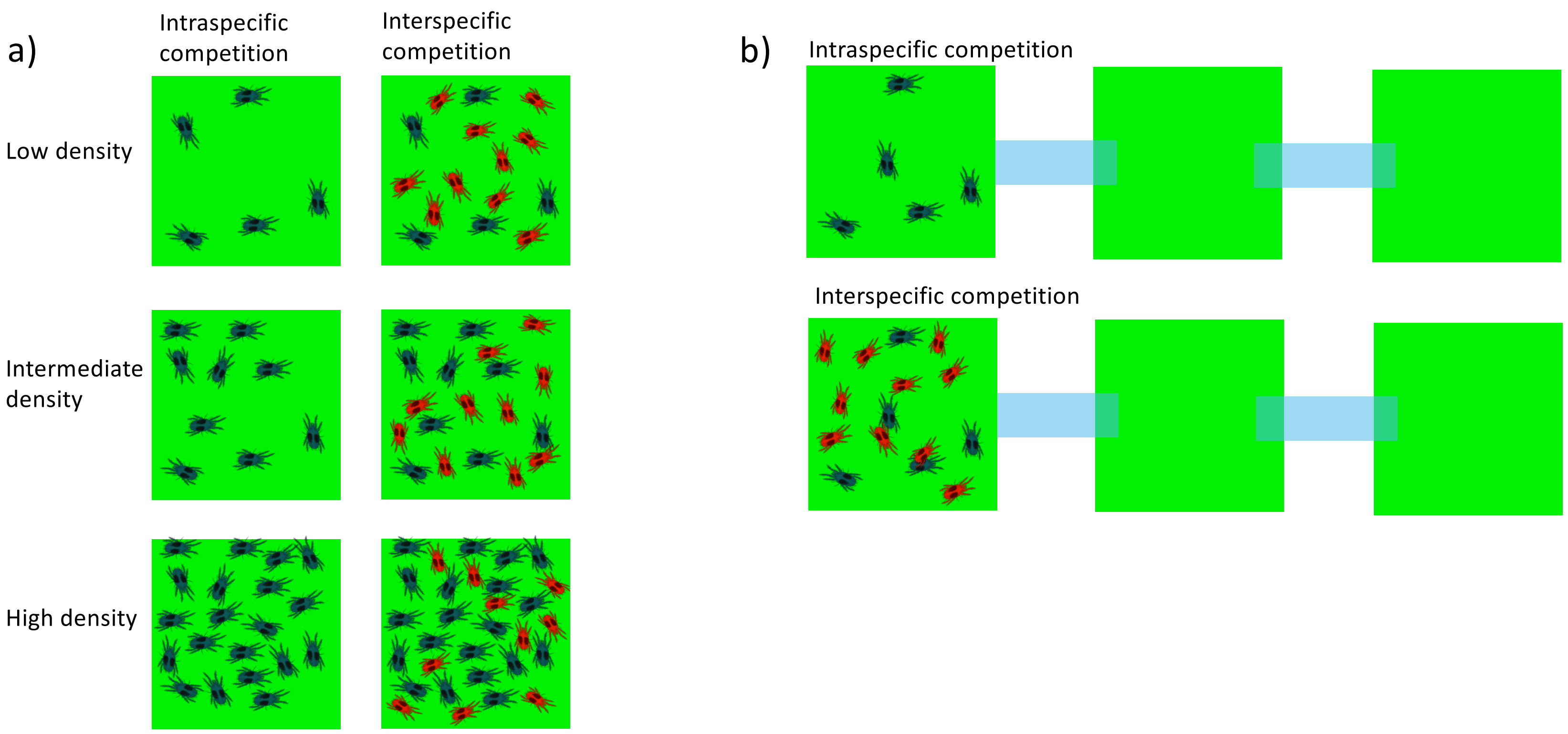

Figure S2: shows mean values (± standard error) of a) virulence (area leaf damage) and b) number of adult daughters at low, intermediate and high density in the presence (red circles) and absence (blue triangles) when traits were measured in the within-host environment.

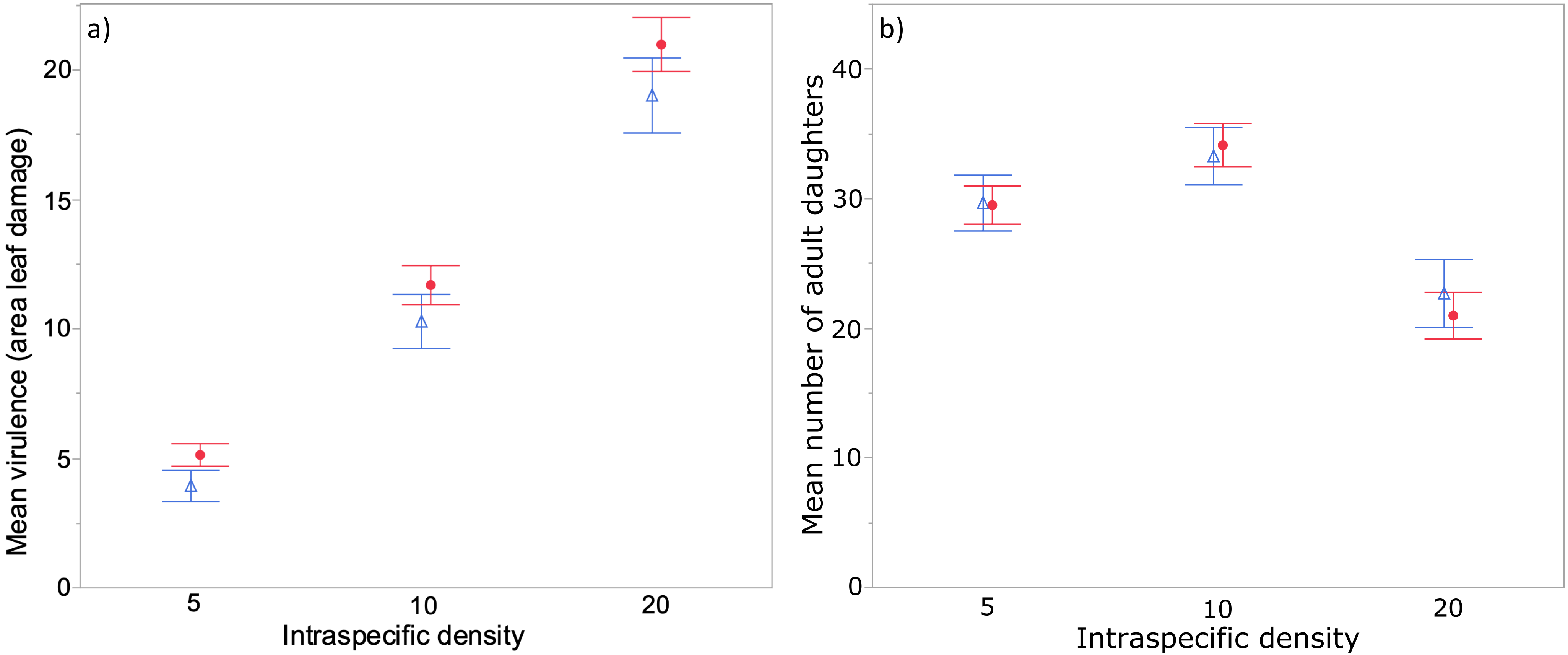

Figure S3: shows the number of mites (± standard error) on a) patch 2 and b) patch 3 through time at each of the intraspecific density treatments in the presence (red circles) or absence (blue) of interspecific competitors

Results: *T. urticae* females colonised host patch 2 approximately 1 day sooner when in the presence of interspecific competitors (Table S4; Figure S3a). The presence of *T. evansi* also increased the maximum proportion of *T. urticae* on reaching host patch 2 (Table S4). The time to reach and the maximum proportion of mites on patch 2 was also higher with increasing *T. urticae* density, but there was no interaction with *T. evansi* competition (non-significant interaction terms; Table S4); the effect of interspecific and intraspecific competition on these traits was additive. The time to reach patch 3 was sooner in the presence of *T. evansi* at low, but not intermediate or high, *T. urticae* densities (Table S4; Figure S3b). The maximum number or proportion of *T. urticae* on patch 3 was unaffected by interspecific competition.

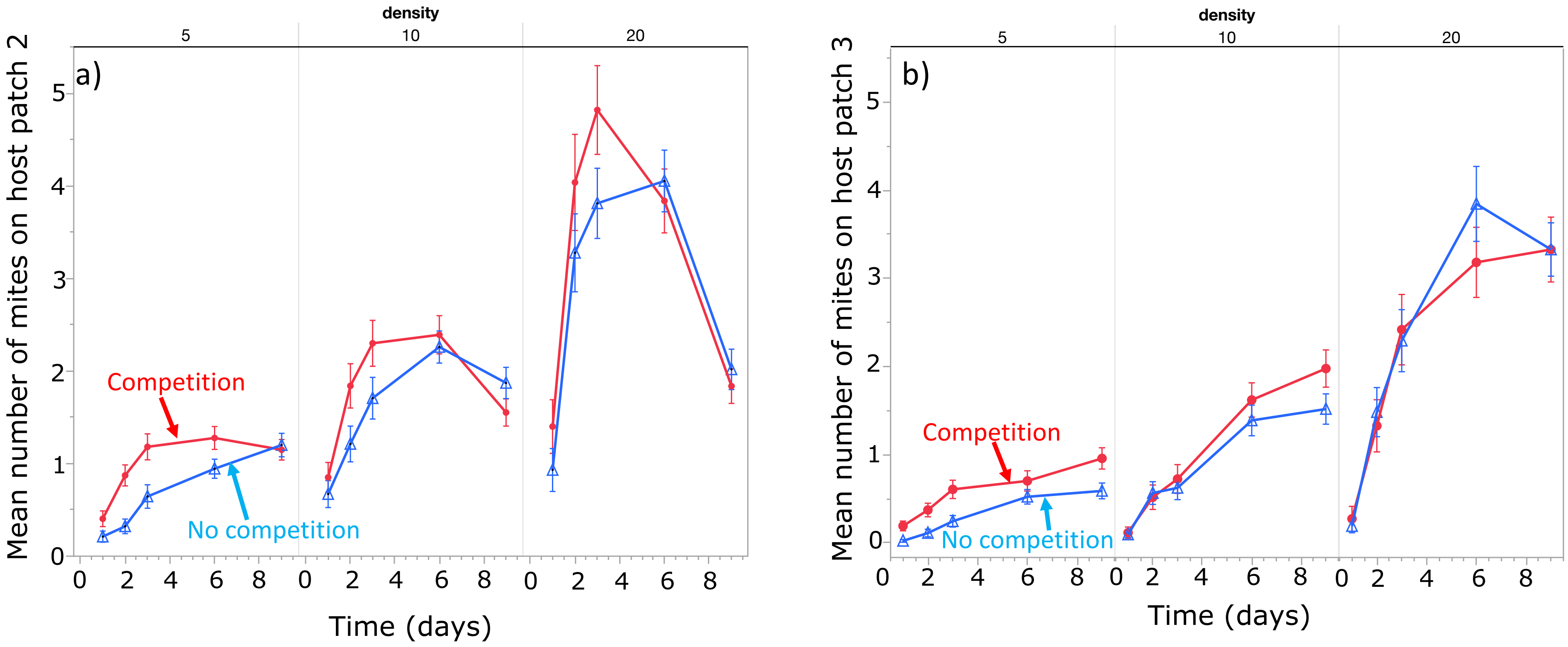

Figure S4: Genetic correlations between a) virulence and day arriving at host patch 2 from host patches with high intraspecific density and b) number of adult daughters and maximum number on host patch 2 at intermediate intraspecific density. Each point is the mean value for an inbred line (± standard error) in the presence (red circles) and absence (blue triangles) of interspecific competition. Note, these figures show standardised vaues of traits with mean of zero and standard deviation of 1.

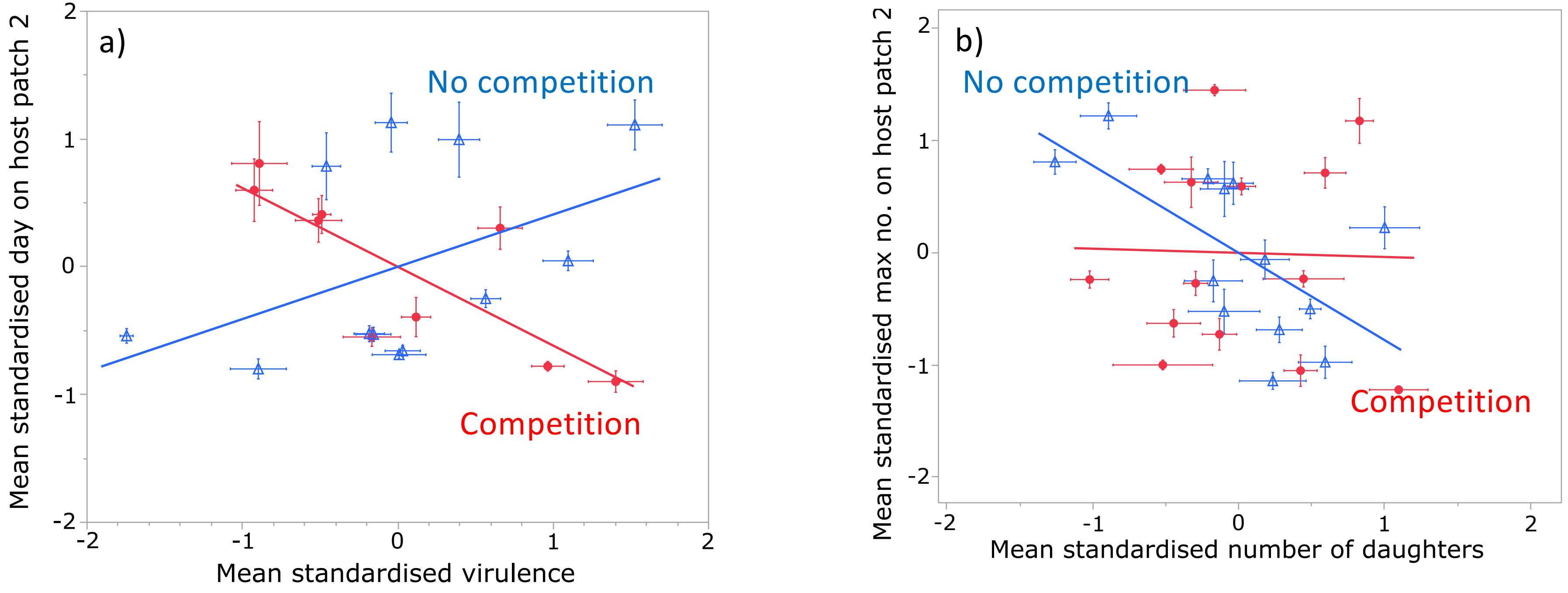

Table S1: Genetic and phenotypic correlations between virulence and number of daughters measured in the within-host environment (on bootstrapped data). Correlations are done separately for each of the different *T. urticae* densities and in the presence or absence of interspecific competition. The genetic correlation ($r_g$) for each pair of traits is presented ± the standard error and the result of the log likelihood test comparing models with and without the genetic correlation. All values of $p < 0.05$ were corrected using Bonferronni (counting 12 tests per pair of traits). Significant correlations are shown in bold.

| | **Type of correlation** | **Density 5** | | **Density 10** | | **Density 20** | |
|---|---|---|---|---|---|---|---|
| | | **No competition** | **Competition** | **No competition** | **Competition** | **No competition** | **Competition** |
| Correlations between virulence and number of adult daughters measured in the within-host environment | Genetic | $r_g = 0.42 \pm 0.24$ SE<br>$\chi^2 = 2.4$, $p = 0.1213$ | $r_g = 0.47 \pm 0.22$ SE<br>$\chi^2 = 3.1$, $p = 0.0783$ | $r_g = 0.28 \pm 0.27$ SE<br>$\chi^2 = 1.00$, $p = 0.3137$ | Model doesn't converge | Model doesn't converge | Model doesn't converge |
| | Phenotypic | **$r_p = 0.24$,**<br>**$p < 0.0012$** | **$r_p = 0.30$,**<br>**$p < 0.0012$** | $r_p = 0.09$<br>$p = 0.1197$ | **$r_p = -0.23$**<br>**$p < 0.0012$** | **$r_p = -0.24$**<br>**$p < 0.0012$** | **$r_p = -0.25$**<br>**$p < 0.0012$** |

Table S2: GLMM investigating the effect of interspecific competitors on total virulence and the total number of offspring in Experiment 1. Mean values for virulence for T. evansi were subtracted from each measure in the competition treatment. The variance components (± standard errors) are presented for the random terms from each minimal model.

| | **Total virulence** | | | **Total number of adult daughters** | | |
|---|---|---|---|---|---|---|
| | **d.f.** | **F ratio** | **P-value** | **d.f.** | **F ratio** | **P-value** |
| Intraspecific density | **1, 648** | **340.03** | **<0.0001** | **1, 651** | **31.90** | **<0.0001** |
| Interspecific competition | 1, 644 | 0.39 | 0.5307 | 1, 650 | 1.23 | 0.2672 |
| ID*IC | 1, 643 | 0.37 | 0.5428 | 1, 643 | 0.06 | 0.8021 |
| Inbred line | 6.27 ± 2.97 | | | 15.87 ± 9.69 | | |
| Block | 24.1 ± 24.38 | | | 39.47 ± 41.29 | | |

Table S3: Results are from a General Linear Mixed Model investigating how competition, *T. urticae* density and time affects the dispersal score. The variance components (± standard errors) are presented for the random terms from each minimal model.

| | Dispersal score | | |
|---|---|---|---|
| | df | F-ratio | P-value |
| Density | 1, 1229 | 15.86 | <0.0001 |
| Interspecific | 1, 450 | 20.82 | <0.0001 |
| Time | 1, 1872 | 691.29 | <0.0001 |
| Density*Interspecific | 1, 451 | 8.76 | 0.0032 |
| Density*Time | 1, 1872 | 0.08 | 0.7742 |
| Time*TIme | 1, 1872 | 151.03 | <0.0001 |
| Density*time*time | 1, 1872 | 22.02 | <0.0001 |
| Line | 0.006 ± 0.003 | | |
| Block | 0.004 ± 0.006 | | |
| Patch | 0.025 ± 0.003 | | |

Table S4: Dispersal related traits in Experiment 2. Results are from General Linear Mixed Models investigating time for *T. urticae* females to reach patches 2 and 3 and the maximum number of adult females on patches 2 and 3. Terms presented are from minimal model for each explanatory variable. The variance components (± standard errors) are presented for the random terms from each minimal model.

| | Time to reach patch 2 | | | Time to reach patch 3 | | | Maximum number of mites on patch 2 | | | Maximum number of mites on patch 3 | | | Maximum proportion of mites on patch 2 | | | Maximum proportion of mites on patch 3 | | |
|---|---|---|---|---|---|---|---|---|---|---|---|---|---|---|---|---|---|---|
| | d.f. | F ratio | P-value | d.f. | F ratio | P-value | d.f. | F ratio | P-value | d.f. | F ratio | P-value | d.f. | F ratio | P-value | d.f. | F ratio | P-value |
| Intraspecific density | **1, 459** | **18.09** | **<0.0001** | **1, 463** | **8.27** | **0.0042** | **1, 466** | **271.65** | **<0.0001** | **1, 466** | **261.14** | **<0.0001** | **1, 461** | **12.03** | **0.0006** | 1, 465 | 0.003 | 0.9535 |
| Interspecific competition | **1, 448** | **6.81** | **0.0094** | **1, 451** | **5.54** | **0.0190** | 1, 453 | 2.68 | 0.1020 | 1, 453 | 1.83 | 0.1768 | **1, 450** | **4.43** | **0.036** | 1, 452 | 1.20 | 0.2751 |
| ID*IC | 1, 449 | 0.30 | 0.5843 | **1, 456** | **5.01** | **0.0258** | 1, 454 | 0.05 | 0.8263 | 1, 454 | 0.04 | 0.8494 | 1, 451 | 0.9862 | 0.3212 | 1, 452 | 0.2135 | 0.6443 |
| Inbred line | 0.79 ± 0.42 | | | 0.49 ± 0.35 | | | 0.09 ± 0.07 | | | 0.09 ± 0.08 | | | 0.002 ± 0.001 | | | 0.002 ± 0.002 | | |
| Block | 0.11 ± 0.20 | | | 0.05 ± 0.01 | | | 0.11 ± 0.18 | | | 0.42 ± 0.62 | | | 0.0009 ± 0.002 | | | 0.005 ± 0.008 | | |

Table S5: Heritability estimates for the number of adult daughters measured in the within-host environment and dispersal related traits in the between-host environment. H2 is calculated as the proportion of the among line variance in the model explained by inbred line. To determine whether inbred line explained a significant amount of the variance for the traits measured, the Akaike's Information Criterion was obtained for models including and excluding inbred line and differences in AIC values > 2 were considered significant (shown in bold).

| | Trait | H2 | AIC with inbred line | AIC without inbred line |
|---|---|---|---|---|
| **Within-host environment** | Number of adult daughters | **0.057** | **3169** | 3186 |
| **Between-host environment** | Time to reach patch 2 | **10** | **2304** | 2318 |
| | Maximum number on patch 2 | 0.24 | 1989 | 1990 |
| | Time to reach patch 3 | **0.03** | **2605** | 2608 |
| | Maximum number on patch 3 | 0.02 | 1987 | 1987 |
| | Dispersal Score | **0.04** | **1423** | 1442 |
